# Replaying the tape of ecology to domesticate wild microbiota

**DOI:** 10.1101/2023.07.07.548163

**Authors:** Alberto Pascual-García, Damian Rivett, Matt L. Jones, Thomas Bell

**Affiliations:** Institute of Integrative Biology, ETH, Zürich, Switzerland; Centro Nacional de Biotecnología, CSIC, Madrid, Spain; Department of Natural Sciences, Faculty of Science and Engineering, Manchester Metropolitan University, Manchester, United Kingdom; Environment and Sustainability Institute, University of Exeter, Penryn, Cornwall, United Kingdom; Imperial College London, Silwood Park, Ascot,United Kingdom

**Author notes:** Equal contribution (Alberto Pascual-García, Damian Rivett).

## Abstract

Humanity has benefited from the domestication of nature and there is an increasing need to predict and control ecosystems. Domesticating bacterial communities would be particularly useful. Bacterial communities play key roles in global biogeochemical cycles, in industry (e.g. sewage treatment, fermented food and drink manufacturing), in agriculture (e.g. by fixing nitrogen and suppressing pathogens), and in human health and animal husbandry. There is therefore great interest in understanding bacterial community dynamics so that they can be controlled and engineered to optimise ecosystem services. We assessed the reproducibility and predictability of bacterial community dynamics by creating a frozen archive of hundreds of naturally-occuring bacterial communities that were repeatedly revived and tracked in a standardised, complex environment. Replicate communities followed reproducible trajectories and the community dynamics could be closely mapped to ecosystem functioning. However, even under standardised conditions, the communities exhibited tipping-points, where a small difference in initial community composition created divergent outcomes. We accurately predicted ecosystem outcomes based on initial bacterial community composition, and identified the conditions under which divergent ecosystem outcomes may be expected. In conclusion, we have shown the feasibility of our approach to reproducibly achieve predictable compositions and functions from wild communities. Nonetheless, the predictability of community trajectories, and therefore their utility in domestication, requires detailed knowledge of rugged compositional landscapes where ecosystem properties are not the inevitable result of prevailing environmental conditions but can be tilted toward different outcomes depending on the initial community composition.

---

Wild bacterial communities are increasingly being used to manipulate ecosystems. Bacterial communities have been transplanted between microbiomes to improve human health [1] and between sewage treatment works to initiate or improve the degradation of organic wastes [2, 3]. They have been inoculated into soil to increase crop yields [4], suppress pathogens [5] and restore wilderness areas following human disturbance [6], and applied to a range of contaminated environments to remove pollutants (bioremediation) [7]. As with approaches that use synthetic bacterial communities assembled from pure cultures, whole-microbiome transplants recognise that multispecies bacterial communities may have more potential than single species [8] because, by acting together, multi-species communities may perform functions with increased capacities [9] and with increased stability due to division of labour [10] and by filling all the available ecological niches [11]. The transplantation approach could be a leap forward in bacterial domestication since it is not restricted to the minority of taxa that can be isolated in pure culture, and bypasses the problem of identifying - from a potentially unlimited number of possible combinations - those consortia that can coexist and function optimally together [12].

The transplantation approach could play a more prominant role in human health, agriculture, bioremediation and other fields if wild communities can be domesticated, but it is unclear whether wild communities are generally suitable for domestication. Requirements for domestication include that wild communities must be stored (e.g. frozen) and propagated, that they must have predictable and reproducible dynamics when they are introduced into novel environments, and that they must provide products and services at predictable and reproducible rates. Unfortunately, the dynamics of multispecies bacterial communities are difficult to predict. Even if individual populations are well-studied, predicting the dynamics of complex communities is challenging because of in-built contingencies; a slight, stochastic increase in the abundance of one population could have cascading impacts that steer the community along a divergent trajectory. Bacterial community dynamics may be broadly predictable in simple environments [13, 14, 15] but usually only at a coarse level of taxonomic resolution [16, 17], likely because taxa that are closely related also occupy similar ecological niches [18]. Bottom-up domestication of bacterial communities has been demonstrated for some synthetic consortia comprising well-characterised or genetically modified strains [19] that in some cases can control proscribed ecosystem properties [20] or confer phenotypes to a host [1, 21]. However, intact wild communities are much more diverse, typically containing thousands of inter-dependent taxa that compete for resources, exchange metabolites, and exhibit sophisticated coordinated behaviours like quorum sensing. Furthermore, high levels of niche overlap among taxa (functional redundancy) can lead to a lottery for community membership and therefore to community dynamics that are governed by chance colonisation order [22].

These observations and ideas make two, apparently contradictory, predictions. The first idea predicts *convergent* community trajectories under standardised environmental conditions. If bacterial communities contain high levels of diversity, natural selection would be expected to rapidly and reproducibly sort the best-adapted taxa, resulting in a single taxonomic and functional outcome. This idea is supported by studies showing a predictable simplification and convergence of communities in standardised conditions [16]. The second idea predicts *divergent* community trajectories that do not reproducibly achieve a certain level of functioning. If many bacteria have overlapping functional capacities, there may be multiple alternative community states that fill the available ecological niches, as implied by studies that have shown taxonomic divergence but functional convergence [23]. This idea is further supported by observations and theory showing that alternative states arise across many study systems, perhaps facilitated by flexible decision-making in bacterial resource acquisition [24]. To address these conflicting ideas, we adapt a famous question from evolutionary biology [25]: does replaying the tape of ecology produce the same outcome? [26].

## Replaying the ecological tape

Here, we ran the tape of ecology 4 times for 275 communities to identify principles of bacterial community domestication. We considered the fate of communities with different initial taxonomic compositions inoculated into a standardised, complex, sterile environment. Each starting community was a naturally-occuring community of heterotrophic bacteria involved in the degradation of beech (Fagus sylvatica) leaf litter in miniature ponds. Leaf litter degradation is an important ecosystem process because increased degradation results in more rapid biogeochemical cycling thereby increasing the productivity of the ecosystem, while lower degradation results in greater carbon storage. Previous experiments have shown that these communities exhibit a strong relationship between degradation rates and the diversity [27] and taxonomic composition [28] of the communities. This system therefore offers a tractable avenue for studying the domestication of wild bacterial communities associated with the provision of a particular ecosystem service. Wild microbiomes were taken from 275 miniature ponds, the bacterial community was separated from co-occurring biota and from the surrounding environmental matrix, and the whole bacterial communities were cryo-preserved. The frozen communities were revived independently four times, re-grown repeatedly in a standardised microcosm containing a sterile beech leaf-based growth medium (see Fig. 1A). We quantified the taxonomic composition of the communities before they were revived from cryopreservation (starting communities) and their composition and functioning at the end of the experiment (final communities), allowing us understand their reproducibility and trajectory and therefore their potential for domestication.

**Figure 1.**
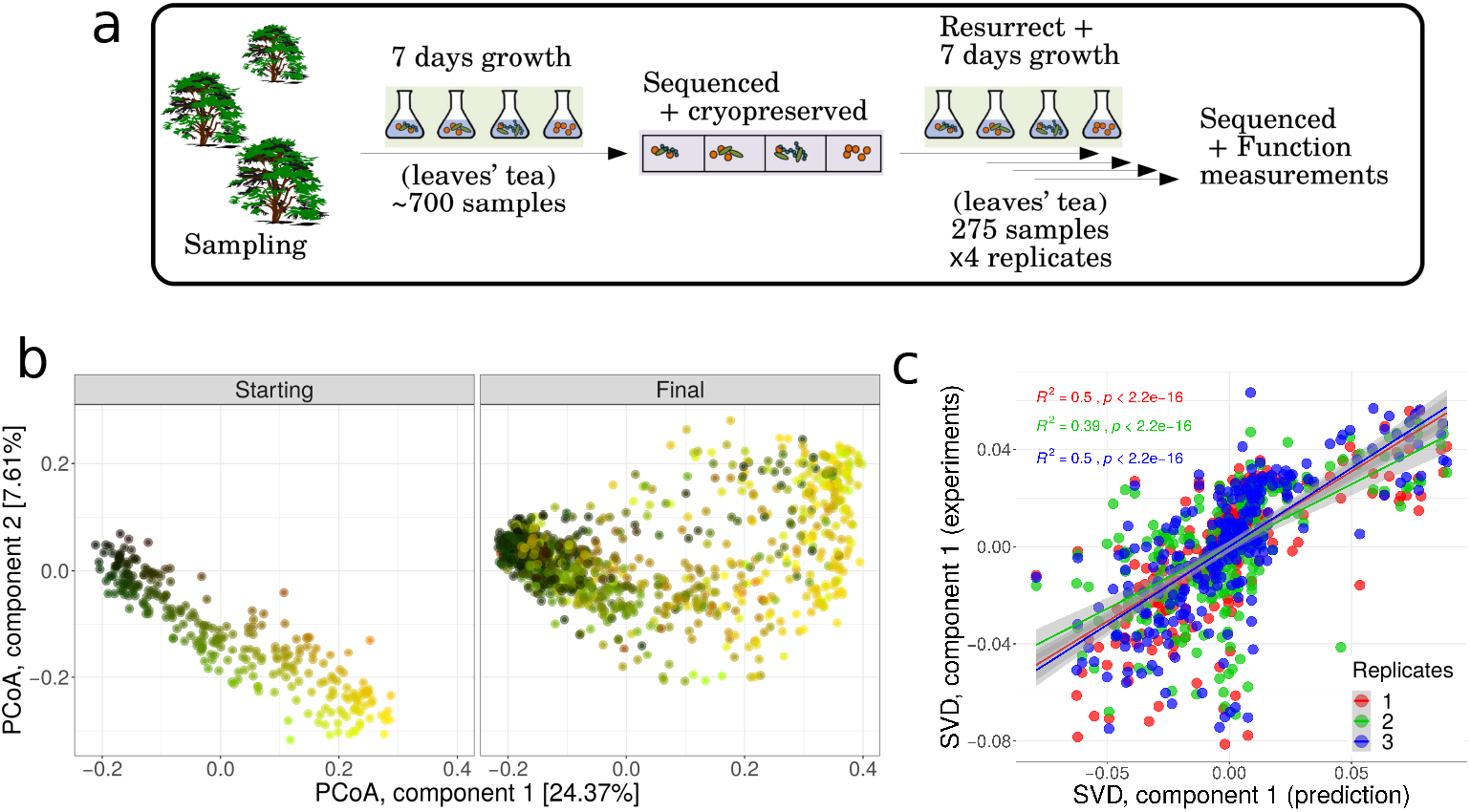
Initial states predict final states. (A) Samples obtained from rainpools were cryo-preserved, and inoculated into a standardised growth medium. Community composition was measured using amplicon sequencing at the start and end of the experiment. (B) Principal coordinates analysis (PCoA) of all communities with colours representing their position in ordination space at the start of the experiment. The left panel shows the starting comunities and the right panel shows the final communities. The third PCoA component is shown in Suppl. Fig. 1.(C) Linear relationship between the Singular Yalue Decomposition (SVD) of transformed starting communities (prediction) and the final communities not used to find the transformation. The transformation accurately predicted the final communities in the other replicates for the first (left panel) SVD components (second SVD component in Suppl. Fig. 2). The Pearson *R*^2^ and significance are indicated for each replicate.

We first identified whether the 4 replicates of each of the 275 communities were clustered at the end of the experiment. We performed a signal-to-noise analysis by quantifying the ANOSIM R statistic [29], which ranges between zero (random groupings) and one (distinct groups). The four replicates produced remarkably non-random groupings (R= 0.716, 275 groups) and there was no difference among the replicates (R= 0.004, 4 groups, see Suppl. Table 1). The reproducibility of the community trajectories was visualised by tracking changes in the abundance of the hundreds of taxa underpinning taxonomic changes using ordination. The ordination revealed that the direction of travel from starting-to final communities was remarkably consistent among communities and replicates (Fig. 1B and Suppl. Fig. 1). To test this idea formally considering the whole of multidimensional space, we asked whether the final composition of one of the replicates could be obtained using a rigid-body transformation (i.e. translation and rotation) of the starting communities. Applying the Kabsch algorithm (see Materials and Methods), we found a small and significant Root Mean Square Deviation between the transformed starting communities and the final replicate selected (0.48, bootstrapped 95% C.I. [0.51, 0.54]). We then asked whether the transformed starting communities predicted the composition of the three replicates that were not used to calculate the transformation by comparing the main components of a singular value decomposition (SVD) of the transformed communities against the SVD of each of the three remaining replicates. We found a strong and significant linear relation (Fig. 1C and Suppl. Fig. 2), confirming that a linear transformation leads to an accurate prediction of the composition of resurrected communities and therefore that groups of taxa collectively move in similar directions in the compositional space.

## Community classes reflect functional differences

At the start of the experiment, the communities were not uniformly distributed across compositional space [30]. To refine our analyses, we therefore performed unsupervised clustering of the communities to identify compositionally-similar sets of communities, similar to the concept of gut microbiome enterotypes [31] (see Materials and Methods). Intuitively, community classes can be regarded as attractors in the compositional landscape, with the ANOSIM R statistic providing a measure of how sharply classes are delimited. We identified an absolute maximum of 17 community classes (R =0.68) and a second maximum with 5 local classes (R = 0. 64). For simplicity and consistency with previous work [30], we analysed the 5 classes (Fig. 2). The starting bacterial communities would have experienced a range of environmental conditions due to site-specific differences in leaf inputs, precipitation, oxygen availability, and many other factors, resulting in a relatively diverse array of community classes 30]. The communities retained signatures of their provenance despite being re-grown twice on the beech leaf media, with communities clustering in compositional space according to their collection location and date (Suppl. Fig. 3).

**Figure 2.**
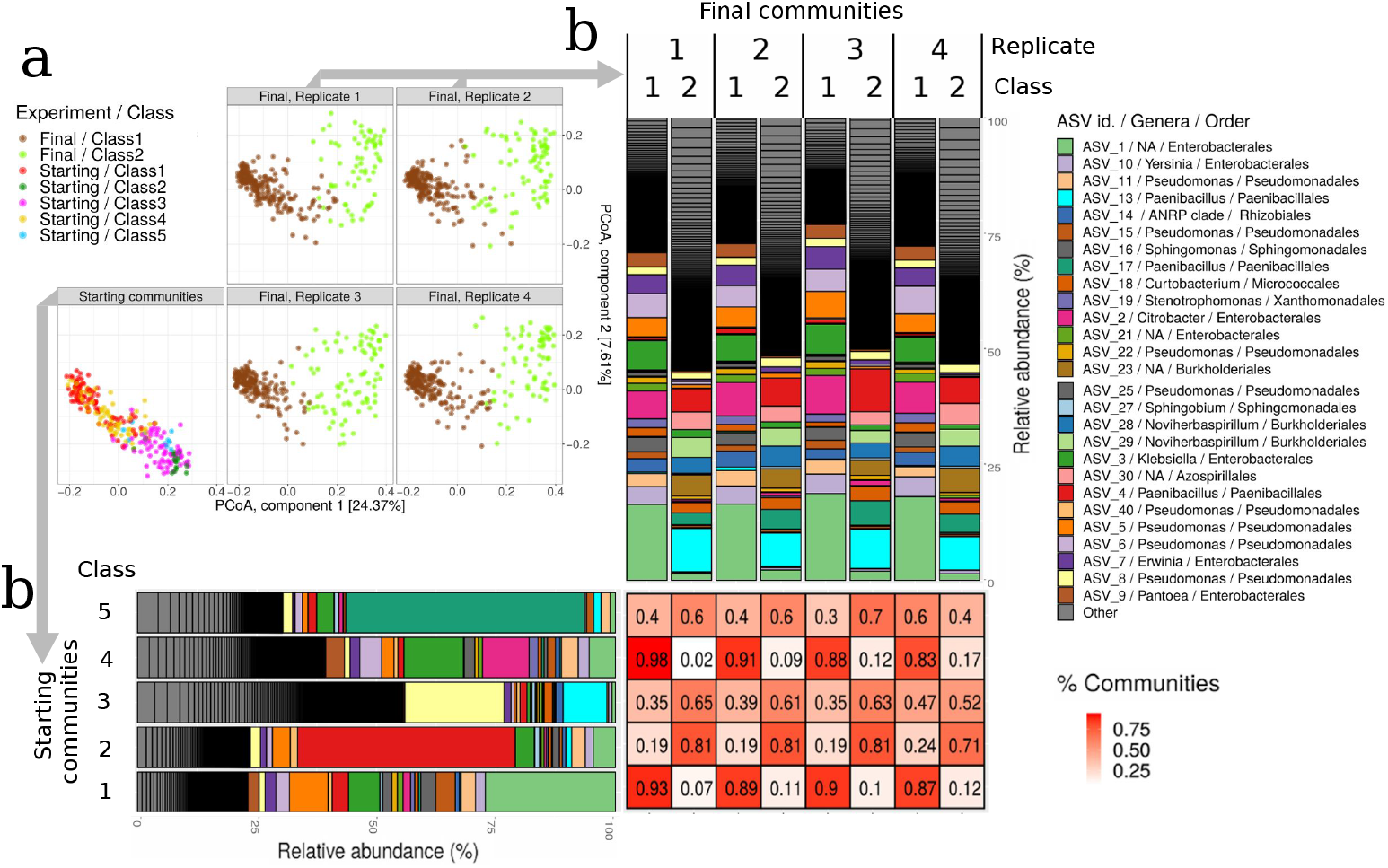
Compositional convergence. (A) Principal coordinates analysis (PCoA) of all communities with colours representing the starting and final community classes. The replicates of the final communities were separated into different panels for clarity to show the reproducible outcome. (B) Bar plots showing the relative abundance of Amplicon Sequence Yariants (ASVs) for starting communities (horizontal bars) and final communities (vertical bars). The communities were divided into the 5 starting community classes and the two final community classes. The final communities were further divided into the 4 replicates. The 20 most abundant ASVs were identified to genus and the remainder were combined into “Other”. Genera that were not resolved are indicated with NA. The matrix indicates the proportion of each starting community class (rows) that resulted in each final community class (columns).

We detected 2 community classes at the conclusion of the experiment (R = 0.78, 2 groups) (Fig. 2 and Suppl. Fig. 4), consistent with the idea that the standardised environmental conditions across the microcosms selected a more limited set of communities that had specific adaptations to the microcosm environment, and demonstrating that separating communities into classes was an economical description of the outcome of the experiment. This conclusion is further illustrated by the observation that 80% of communities had all four replicates ending in the same final class, and only 2.5% of the communities had replicates evenly split into the two final community classes. The outcome of each community was therefore highly dependent on the starting community: once the initial composition was known, its fate was ordained with only minor deviations.

Future domesticators of wild bacterial communities are likely to be less interested in predicting the precise composition of domesticated communities, and more interested in predicting the ecosystem services that will be extracted. We therefore explored ways of mapping functional data onto the community trajectories by inferring the functional profiles of the communities from their composition, and by direct functional measurements. We identified associations between the 2 final community classes and the functional performance of the communities by first imputing the metagenomes of these communities from the 16S rRNA sequencing data, using PiCRUST to categorise genes by function using the KEGG database. This analysis suggested that the 2 community classes were associated with distinct modes of nutrient uptake. In Final Class 1, which contained the majority of final communities, there were a higher proportion of genes related to rapid nutrient uptake (e.g. transporters, phospho-transferase system), transcription factors, and fructose and mannose metabolism (Fig. 3A, and Suppl. Figs. 5 and 6). There was also a higher proportion of genes associated with lipo-polysaccharide metabolism and membrane and intracellular structural proteins, which are characteristic of Gram negative bacteria such as *Serratia* and *Pseudomonas* species that were dominant in Final Class 1 (Fig. 3A). By contrast, Final Class 2 communities had a higher proportion of genes related to oxidative phosphorylation and related pathways to acquisition of acetyl-coA (e.g. degradation of amino acids, fatty acids metabolism and propanoate metabolism). Final Class 2 also retains pathways that were more abundant in starting classes 2 and 5. These pathways include genes related to chemotaxis and motility and some expected in scarce environments such as sporulation genes (Suppl. Fig. 7). Since these starting classes had the lowest number of communities and Final Class 2 also hosts less communities than Final Class 1 (Suppl. Table 1), this suggested that the majority of communities gravitated toward improved leaf litter uptake, while a minority may be relics of communities that were poorly adapted to the beech leaf culture media or became self-limiting (e.g. due to build up of waste products) [30].

**Figure 3.**
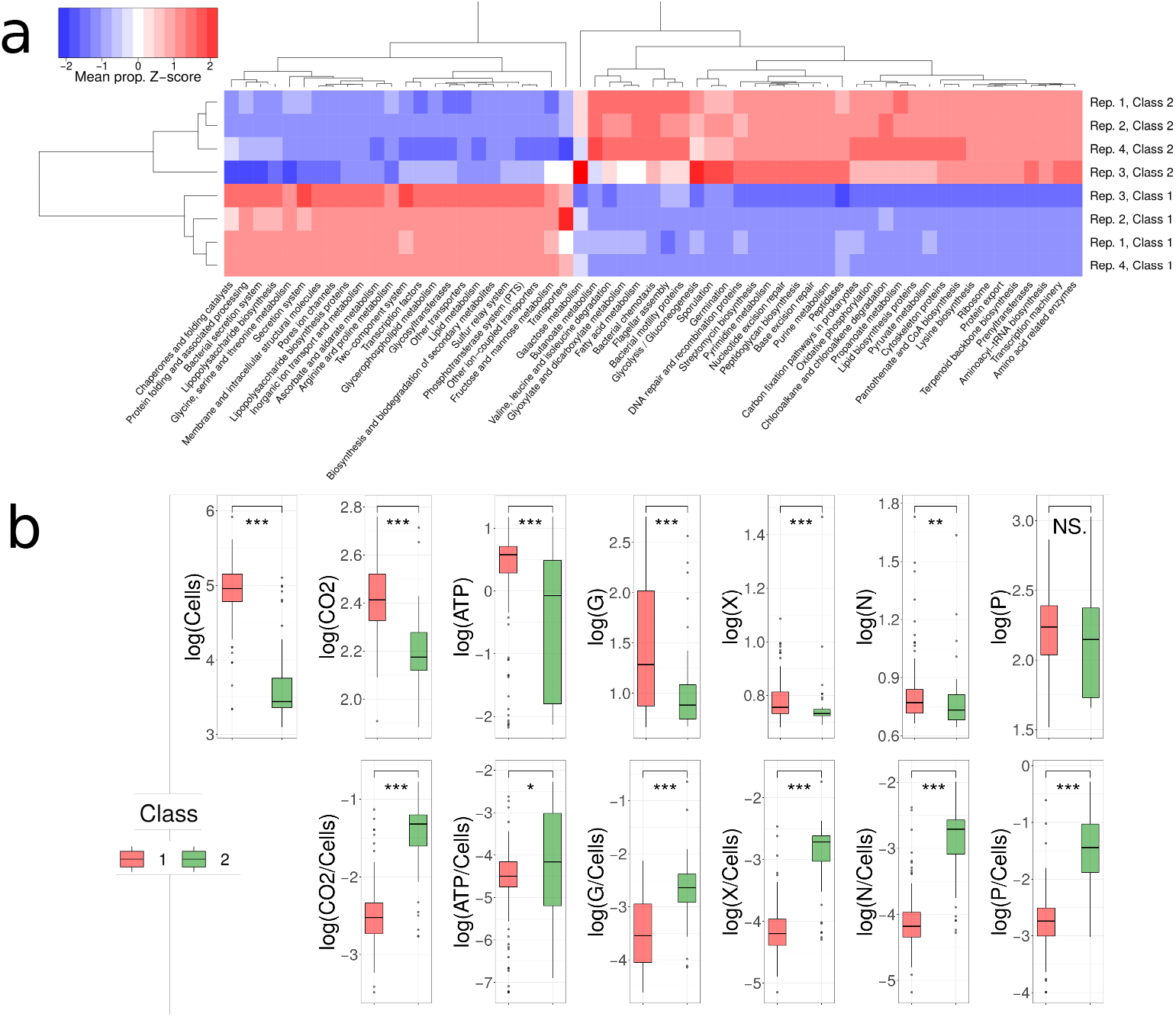
Functional convergence. (A) Z-score of the mean proportion of genes clustered in KEGG metabolic pathways found in final community classes. The scaling of the Z-score was computed for each pathway (i.e. scaled by columns). Only pathways showing a significant difference between at least two classes are shown. (B) The functional measurements were significantly different betweem the two final community classes. All replicates were merged. G: *β*− glucosidase; X: xylosidase; N: *β*-chitinase; P: phosphatase. Wilcoxon test, significance p-value: *** = 0.001, ** = 0.01, * = 0.05, NS not significant.

We further validated these metagenomic predictions using direct measurements of functional performance of the final communities by measuring the degradation rate of four substrates along with the whole-community metabolic activity, respiration rate, and cell numbers. Final Class 1 communities had higher degradation rates and enzymatic activity, while Final Class 2 communities had a higher per-cell investment but lower cell numbers, implying that they were growing inefficiently (Fig. 3B). We speculate that this was due to higher metabolic costs in this environment for tasks not associated with cell growth and division such as exo-enzymatic investment, and that these communities may be adapted to environments with fewer or more recalcitrant resources. Regardless of the underlying mechanism, both the metagenomic profiles and the functional experiments demonstrate that the the final community classes are not just alternative taxonomic variants providing the same functional outcome, but that the divergent community classes translate into significant functional differences. The two final classes are therefore unlikely to result from neutral community drift.

## From tape to landscape

Aggregating the communities into community classes allowed us to quantify the reproducibility of community trajectories by identifying whether the fate of communities originating from the same class was fixed. We found that, although individual communities had remarkably reproducible trajectories, the trajectory of each community was contingent on the initial community class. We illustrate this contingency by showing the fate of each of the initial 5 starting communities classes. Starting Class 1 and Class 4 communities consistently converged toward final class 1 communities, with approximately 90% of the starting communities originating from those classes (Fig. 2). A hypothetical study that used one or a few communities that were within either of those classes would have observed a single outcome, consistent with the idea that communities converge and simplify under standard conditions. Other communities were less predictable in their trajectories at the class level, with outcomes for communities in Starting Classes 2, 3, and 5 more evenly split between Final Class 1 (∼20-50%) and Final Class 2 (∼50-80%) final communities (Fig. 2). A study that used one or a few of the communities from starting community classes 2, 3, or 5 would therefore likely have observed the formation of alternative compositionally and functionally divergent outcomes.

Conceptually, the experimental communities can be visualised as traversing a compositional and functional landscape (Fig. 4A), with communities starting from compositions set by the their native environmental conditions, and where community stability across the landscape is determined by the new environmental conditions. Stable communities would be those that are resistant to change and would return to their original composition if they are perturbed, while unstable communities would be less resistant to change. Communities would rapidly traverse compositional space containing unstable compositions and would spend more time in neighbourhoods where the communities were more stable. If there was a single composition in the landscape that was stable, any starting community composition would generally converge on this single ‘attractor’ [30]. Communities located near saddle points or on ridges of the landscape would have a more unpredictable outcome since initial small changes to the composition would tilt the communities toward alternative peaks. If all compositions were equally unstable, the community dynamics would drift from one unstable composition to another, resulting in a random walk through compositional space with divergent, unpredictable outcomes among communities and among replicates. The topography of the landscape determines the predictability and repeatability of the community dynamics and therefore provide information about whether the communities can be domesticated (Suppl. Fig. 8).

**Figure 4:**
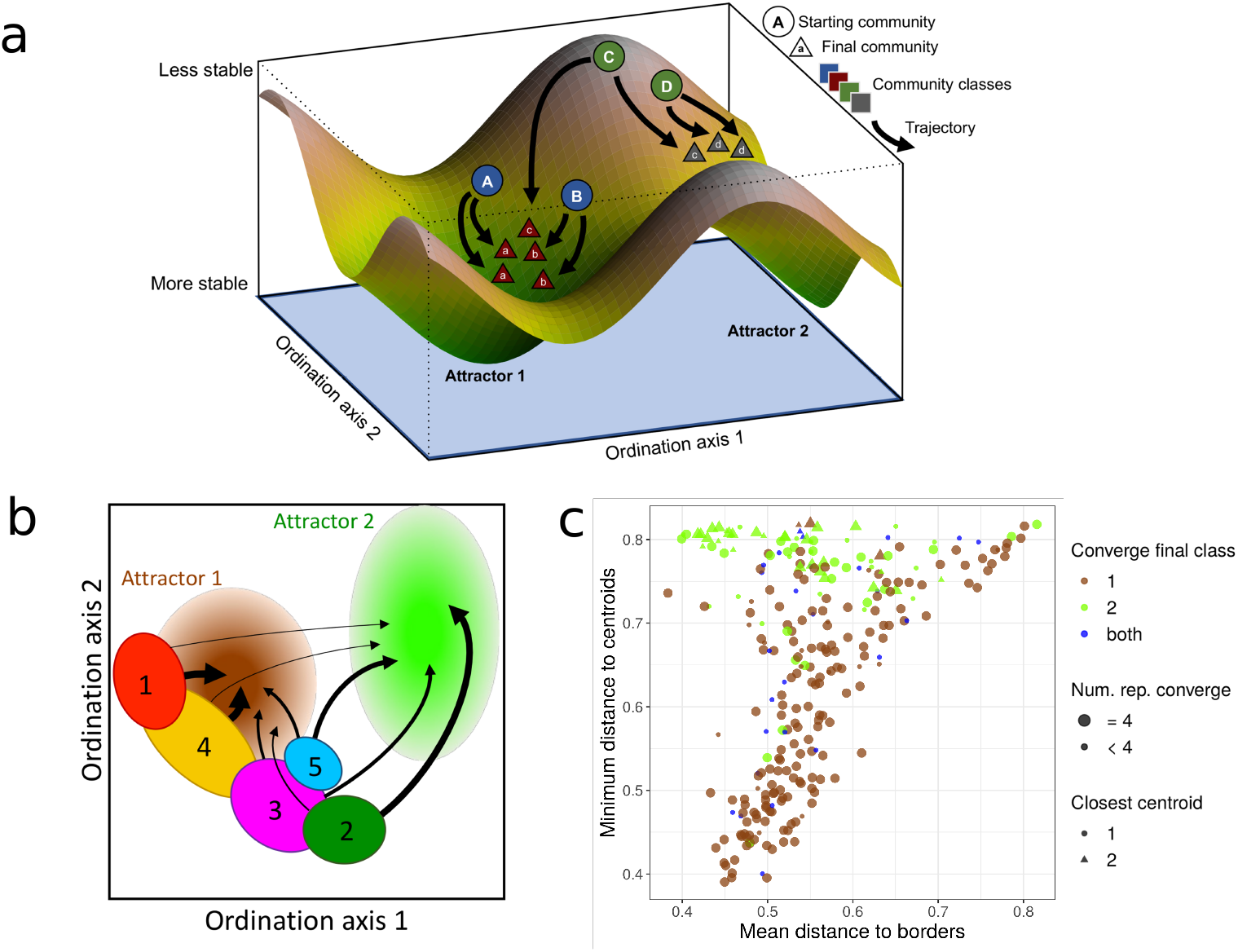
Community trajectories are influenced by the topography of the compositional landcape. (A) Illustration of the compositional landscape from the perspective of a set of starting communities entering a new environment. Two attractors are shown, with circles representing the starting communities, triangles representing the final communities, shape colours indicating community classes, and arrows indicating community trajectories. (B) Illustration of the location (in ordination space) and trajectory of each starting class used in the experiment. Ovals indicate approximate locations of the numbered starting classes, arrows indicate trajectories with thicker arrows indicating more consistent outcomes, and pale circles indicating putative attractors. An equivalent representation with real data is provided in Suppl. Fig. 9 (C) For each starting community, we computed the mean distance to the centroid of the final community classes and the minimum distance to the borders (closest community in the class). We observed that starting communities that were distant from both borders converged to Final Class 2 particularly when they were closer to its centroid.

The experimental results are consistent with a rugged landscape containing at least two attractors toward which the composition tended to gravitate (i.e. Fig. 4B). Starting community Classes 1 and 4 lie along the flank of the attractor close to final community class 1, resulting in convergence to a single outcome (Fig. 4B and Suppl. Fig. 9). By contrast, Starting Classes 2, 3 and 5 sit on a ridge that allows them to diverge to Final Class 1 or 2 (Fig. 4B and Suppl. Fig. 9). To quantify this idea we computed the distance of each starting community to the centroids and borders of the final attractors (Fig. 4C). Communities that were further away from both centroids tended to converge to Final Class 2 particularly those starting communities that were closer to both its centroid and border. This suggests that the final community class 2 attractor is less steep than the class 1 attractor or was generally less accessible from the starting community compositions. Starting communities needed to be far from the final community class 1 attractor to avoid its pull, and would only converge to Final Class 2 if they were sufficiently distant. This was further confirmed when we computed the mean distance between starting and final classes, with Starting Classes 1 and 4 (the largest set of communities) having a high similarity with Final Class 1, suggesting that these two starting classes were already orbiting an attractor that, after the second round of growth, consolidated into a single large and robust attractor (Suppl. Fig. 10). By contrast, Starting Classes 2, 3 and 5 were more similar to Final Class 2, but their mean similarity was much lower, suggesting a substantial compositional transformation of the communities falling into this attractor (Suppl. Fig. 10). Selecting communities for domestication would therefore require detailed knowledge of rugged compositional landscapes or would be at risk of diverging to undesirable compositional and functional outcomes.

## Conclusions

The experimental results demonstrate the reproducibility of community dynamics, a central requirement for understanding how microbial communities assemble and for establishing their utility as agents for altering ecosystems. By preserving and reviving hundreds of communities we were able to expand on this simple discovery. First, there was a strong historical contingency in the community trajectories, with naturally-occuring compositional classes from the field predicting the compositional classes that communities gravitated towards in the laboratory. Second, these broad compositional classes towards which communities gravitated also reflected the main axes of functional performance, both in terms of their functional capacity (metagenomes) and their delivery of ecosystem processes (degradation rates and activity). Third, while the community dynamics were highly predictable under standardised initial conditions, the trajectory depended on the initial community composition. Communities did not simplify to a single outcome, but individual communities may have different (predictable) trajectories. Together, the results show the feasibility of pushing ecosystems toward alternative compositional and functional outcomes by minor alterations to the community composition, but that the capacity to do this depends on the topography of the compositional landscape.

## Materials and Methods

### Laboratory methods

We sampled 753 rainwater pools from the buttressing of beech trees (*Fagus sylvatica*) during August 2013 to April 2014 [28]. The pools were stirred thoroughly to obtain an unbiased sample of the whole community and we collected 1 ml of water and sediment. The samples were diluted 1:4 in sterile phosphate buffered saline (PBS, pH 7.0, Sigma-Aldrich) and filtered (pore size 20-22 µm, Whatman 4 filter paper) to remove debris. The filtrate containing the communities was inoculated into 5 ml sterile beech leaf medium (50 g dried beech leaves autoclaved in 500 ml PBS, filtered, diluted 32-fold in PBS, amended with 200 µg ml−1 cyclohexamide (Sigma-Aldrich) to inhibit fungi). Each community was incubated at 22°C under static conditions for 1 week to allow communities to reach stationary phase. A sample was collected to characterise the (starting) community composition (16S rRNA sequencing, see Sequencing methods) and communities were stored at -80°C after addition of glycerol, which acts as a cryoprotectant (final concentration 30% v/v glycerol, 0.85% w/v NaCl). Further details of the experimental methods on starting communities are provided elsewhere 28], which also includes documentation of how cryopreservation impacted the communities.

The experiment to revive and grow final communities was conducted in microcosms (1.2-ml-deep 9 -well plates) containing 840 µl sterile beech leaf medium inoculated with 40 µl of each revived community (∼20,000 cells). Each community was revived indpendently four times, and 275 communities were assayed, yielding a total of 1,100 microcosms. The microcosms were incubated under static conditions at 22 ° C for 7 days, after which time the communities were characterised for their taxonomic composition (see Sequencing methods) and measured ecosystem functioning.

We measured bacterial activity, growth, and rates of substrate degradation in the microcosms. The measurements were generally taken at the conclusion of the experiment except for respiration, which was measured cumulatively throughout the experiment. Final bacterial counts were obtained by staining the cells with thiazole orange (42 nM, Sigma-Aldrich) followed by obtaining absolute counts using a C6 Accuri flow cytometer (size threshold of 8,000 forward scatter height (FSC-H)), with cells gated on the side scatter area (SSC-A) and fluorescence channel 1 (FL1-A) (533/30) channels. We used a threshold of 800 fluorescence units to distinguish cells from detritus. Bacterial respiration was measured using the MicroResp CO2 detection system (www.microresp.com) according to the manufacturer instructions, with absorbance readings converted to CO2 using a standard curve [28]. Respiration measurements were taken as the cumulative respiration of the whole community over the 7-day incubation period. Potential for metabolic activity was measured as the adenosine triphosphate (ATP) concentration within the community, measured using a Biotek Synergy 2 multimode plate reader and the BacTitr-Glo Cell Viability assay (Promega). There was a linear relationship between concentration and luminescence (R2 =0.998), which we used to convert luminescence to nM ATP. We measured the breakdown of substrates labelled with 4-methylumbelliferone (MUB). Samples were amended with 40 µM of the substrates (100 µl total volume) and incubated in the dark under the same conditions as the microcosms (static, 22 ° C) for 0 minutes. After the incubation, 10 µl of 1 M NaOH was added and the fluorescence measured over four minutes with the maximum value recorded. Fluorescent values were converted to nM MUB after establishing a linear relationship between MUB concentration and fluorescence (R2= 0.996) and using negative controls to account for any auto-fluorescence in the medium. We selected substrates that were ecologically relevant to this ecosystem including xylosidase (cleaves the labile substrate xylose, a monomer prevalent in hemicellulose), β-chitinase (breaks down chitin, the main component of arthropod exoskeletons and fungal cell walls), β-glucosidase (breaks down cellulose, the structural component of plants) and phosphatase (breaks down organic monoesters for the mineralization and acquisition of phosphorus).

### Sequencing methods

We characterised the composition of each initial community and of the 4 replicate final communities on the Illumina MiSeq platform by Molecular Research DNA (www.mrdnalab.com). The V4 region of the 16S ribosomal RNA gene was amplified, using primers 515f/806r with a barcoded forward primer. The sequencing effort (15000 reads per sample) was similar to the number of cells used to initiate the microcosms (∼20000 cells), so we assumed the communities were almost fully characterized. 16S rRNA amplicon sequence data was processed using the UNIX and R (v4.2.1) programming languages [32]. Briefly, demultiplexed sequence files obtained from the sequencing facility were processed using DADA2 pipeline in R [33, 34] version 1.14 to produce a bacterial amplicon sequence variant (ASVs: [35]) abundance table. The quality profiles of the reads were filtered and trimmed (using the function dada2::filterAndTrim); truncating sequences to 240bp (option trunLen = 240), removing reads with a quality score less than 11 (option truncQ = 11), discarding reads with ambiguous bases (Ns; option maxN = 0) or with more than one expected error (Ns; maxEE option = 1), and removing reads that matched the phiX genome (option rm.phix =TRUE). Error rates were then learned using dada2::learnErrors, before sample inference (ASV inference) using the main dada2::dada function to create an ASV abundance table. Chimeras were removed via the dada2::removeBimeraDenovo function using the consensus method (option method = ‘consensus’) before taxonomic assignment of the ASVs to the species level using dada2::assignTaxonomy, aligning to the SILVA v138 SSU Ref NR 99 database [36, 37, 38]. This process identified 21083 ASVs across all samples. After inference of ASVs from the sequence data in this way, we applied additional, custom quality-filtering procedures - namely, removing ASVs with fewer than 100 reads across samples (reducing the number of ASVs to 5834) and removing samples with fewer than 10000 sequences (reducing the number of ASVs to 1209). This resulted in the final ASV table of 1209 ASV abundances across all of samples from days 0 and 7.

### Visual representation of the communities

ASV tables were rarefied to 10K reads for visualization purposes only. Bar-plot representations were created with phyloseq [39] and dimensionality reduction performed by computing all-against-all communities dissimilarities with Jensen-Shannon divergence (*D*_JSD_) [40], followed by a Principal Coordinates Analysis (R function dudi.pco, package ADE4).

### Determination of communities classes

Following previous work, we determined communities classes by computing all-against-all *D*_JSD_ and performing a Partition Around Medoids clustering, which requires as input the number of output communities *k*. To find the optimal clustering, we ran the method for a broad range of *k* values and computed the Calinski-Harabasz index (*CH*) that quantifies the quality of the classification. The optimal classification results from choosing *k*_opt_ = arg max_*k*_(*CH*), shown in Suppl. Figs 4. This procedure was followed for each subset of data independently (after quality-filtering sequencing procedures, 658 samples for starting communities and 275 samples for each replicate of final communities).

### Optimal community superposition

We superimposed the starting communities on one of the replicates of the final communities using a Kabsch algorithm (adapting the implementation available in the PDBencode R, URL: https://github.com/Fraternalilab/PDBencode), which applies the Singular Value Decomposition (SYD) of the cross-covariance matrix of the relative abundances of both datasets and then seeks an optimal transformation. To obtain an estimation of the quality of the superposition, we repeated the computation after bootstrapping the ASV table 50 times to obtain a confidence interval. Finally, we further evaluated the prediction by representing the first component of the SVD of the transformed starting communities against those of the three remaining replicates of the final communities.

### Community classes and the compositional landscape

To evaluate the significance of the community classes that we identified, and to confront our categorisation with other potential groupings, we computed the ANOSIM metric (R package vegan [41]) assessing the significance with permutation tests (10^3^ permutations). To illustrate the fate of starting communities with respect to the final classes we first computed the centroid of each final class. For each set of communities belonging to a given class, we generated a new set by sampling 10000 reads with the probabilities given by the relative abundances of each sample. The centroid was defined as the median value of each ASV across the resampled set, whose relative abundance was considered for downstream computations. Second, we computed, for each starting community, *D*_JSD_ against all final communities of one of the replicates and its centroids. To generate Fig. 4D we identified the minimum distance to each class. Similar results were obtained regardless of the replicate that was selected.

### Metagenomic predictions

Metagenomic predictions were performed using PiCRUST v2.4.2 [42], which was also used to compute quality controls. Eleven ASVs had poor alignments and were removed from downstream analysis. The nsti score assessing the quality of the prediction was 0.042, indicating that the predictions were of high quality. Predictions were aggregated considering both the KEGG pathway hierarchy and BRITE annotations [43]. For each community class, the proportion of each pathway was averaged across all samples in that class. The difference in the mean proportions was then computed between each pair of classes, and those pathways showing a significant difference in at least one comparison were retained (Welch test corrected for multiple testing, difference in mean proportions larger than 0.05 and corrected *p* − *val* < 0.001, see Suppl. Figs. 5 and for examples). The mean proportion of the pathways selected were represented in a heatmap (Fig. 3A) and rescaled computing a Z-score to highlight those over-(or under-) represented in each class. Rows and columns were clustered with an average linkage agglomerative clustering using an Euclidean distance (default method in heatmap.2, R package gplots [44]).

## Acknowledgements

We thank Lara Durán-Trío for useful discussions.

## Data availability

Sequences associated with this study are deposited at NCBI under BioProject accession number PRJNA989519. This project contains the 16S rRNA amplicon sequencing data associated with each of the communities at day 0 (SUB13586664), as well as at day 7 for the four replicate growth experiments (SUB13586665-8). Full details to reproduce results can be found in the repository of the code provided.

## Code availability

Code used for all the analysis presented in the manuscript was deposited in GitHub with the URL: https://github.com/apascualgarcia/ReplayEcology.

