## Supplementary Materials for "Replaying the tape of ecology to domesticate wild microbiota"

### Supplementary Materials
### Replaying the tape of ecology to domesticate wild microbiomes

Alberto Pascual-García[1,2,♠], Damian Rivett[3,♠], Matt L. Jones[4], Thomas Bell[5,*]

July 8, 2023

(1) Institute of Integrative Biology, ETH, Zürich, Switzerland
(2) Present address: Centro Nacional de Biotecnología, CSIC, Madrid, Spain
(3) Department of Natural Sciences, Faculty of Science and Engineering, Manchester Metropolitan University, Manchester, United Kingdom.
(4) Environment and Sustainability Institute, University of Exeter, Penryn, Cornwall, United Kingdom.
(5) Imperial College London, Silwood Park, Ascot,United Kingdom.
(♠) Equal contribution

### Supplementary Figures

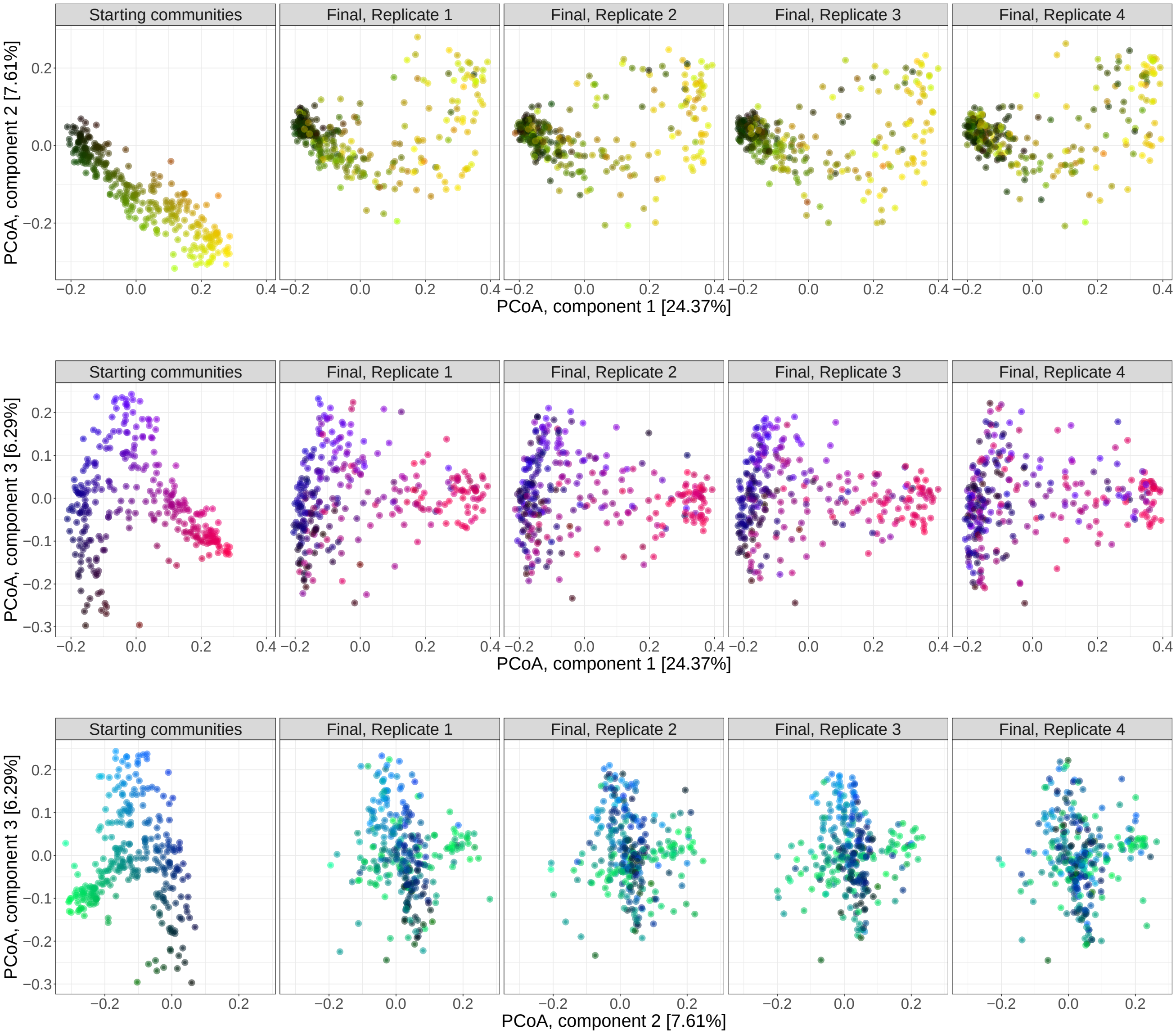

Figure 1: **Illustration of trajectories.** PCoA performed with the Jensen-Shannon divergence and projected on coordinates 1 and 2 (top), 1 and 3 (middle), and 2 and 3 (bottom), with the communities coloured by their starting position in the ordination space. Final replicates were split in different boxes for clarity.

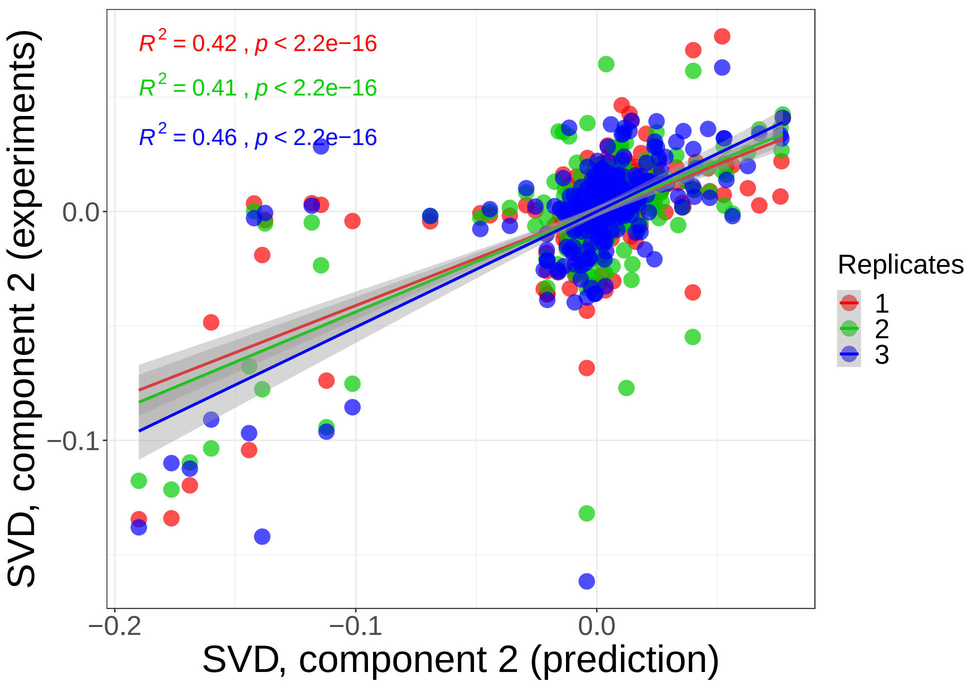

Figure 2: **Predicted final composition** of starting communities after rigid-body transformation against the composition of the replicates not used to find the transformation. The second Singular Value Decomposition component is used for the comparison.

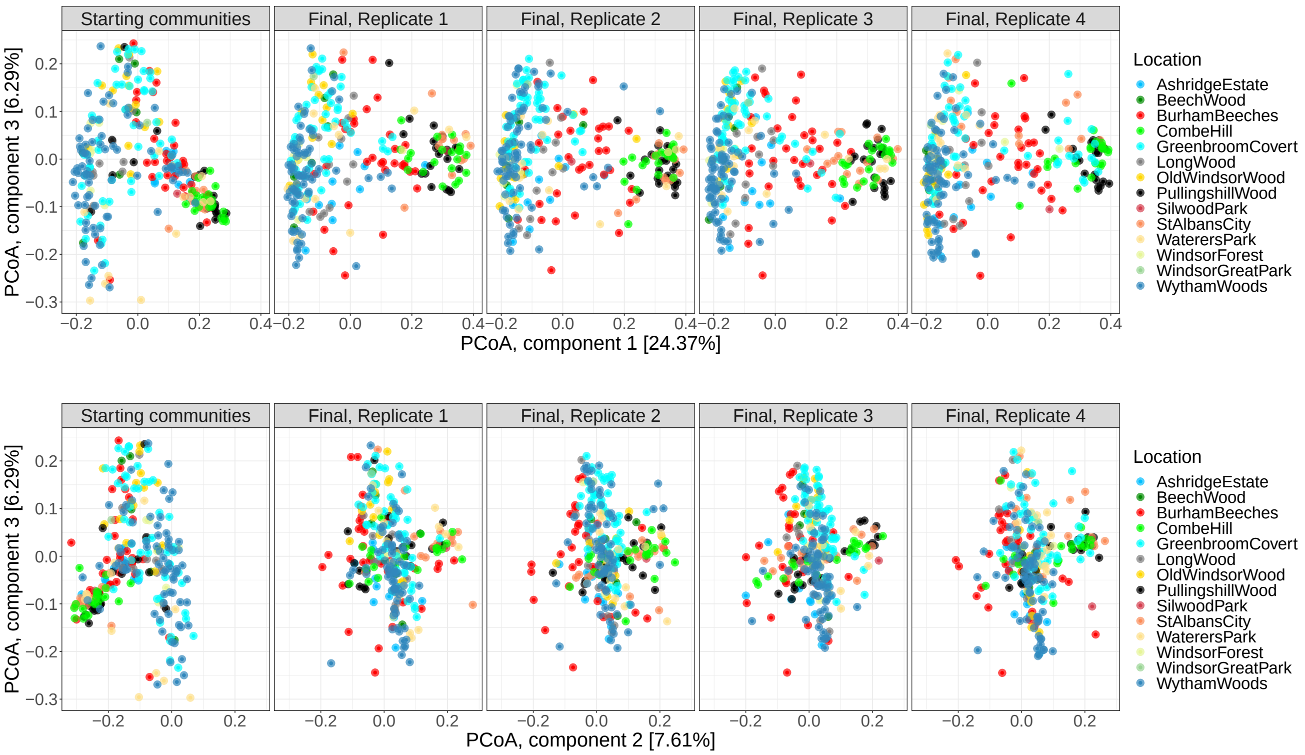

Figure 3: **History of communities.** PCoA performed with the Jensen-Shannon divergence and projected on coordinates 1 and 2 (top) and 1 and 3 (bottom), with the communities coloured by their location. Final replicates were split in different boxes for clarity.

### Supplementary Text

#### Identification of classes and statistical significance

As indicated in the Main Text (see Methods) to identify the number of classes we performed a Partition Around Medoids clustering, which requires as an input the number of clusters $k$. To identify the optimal clustering, we computed the Calinski-Harabasz ($CH$) index and chose the classification having the maximum value of this quantity, shown in Suppl. Fig. 4. In previous work, we followed a similar procedure but worked with just the starting communities and used Operational Taxonomix Units clustered at 97% sequence identity (see [1] for details), which resulted in six clusters (community classes). In this work, we found that working with Amplicon Sequence Variants the number of classes increased to 17 (see Suppl. Fig. 4, left) with a second maximum when the number of clusters was 6. For consistency with previous work we chose this second maximum as our reference classification, and we confirmed that both classifications were qualitative similar, allowing us to interpret our results at the light of previous findings. Only 5 classes were represented in the subset of starting communities that were resurrected in this study. For the final communities, the $CH$ index becomes more skewed, dropping precipitously for any clustering beyond $k = 2$, suggesting that the compositional landscape was simplified with respect to the starting landscape (see Suppl. Fig. 4, right). Finally, the significance of the classes was evaluated by computing the ANOSIM metric, and confronted with other potential groupings, shown in Suppl. Table 1.

| Dataset | Groups | ANOSIM |
|---|---|---|
| All | Starting vs Final | 0.029 |
| All | Parent+Childs | 0.656 |
| Starting | Classes | 0.643 |
| Final | Replicates | 0.004 |
| Final | Classes | 0.780 |
| Final | Same parent | 0.716 |
| Replicate 1 | Classes | 0.707 |
| Replicate 2 | Classes | 0.793 |
| Replicate 3 | Classes | 0.784 |
| Replicate 4 | Classes | 0.839 |

| **Starting communities** | |
|---|---|
| **Class** | **Number** |
| 1 | 30 |
| 2 | 35 |
| 3 | 121 |
| 4 | 39 |
| 5 | 29 |

| **Final communities** | | |
|---|---|---|
| **Replica** | **Class** | **Number** |
| 1 | 1 | 187 |
| 1 | 2 | 67 |
| 2 | 1 | 180 |
| 2 | 2 | 74 |
| 3 | 1 | 171 |
| 3 | 2 | 83 |
| 4 | 1 | 184 |
| 4 | 2 | 70 |

Table 1: **Classes statistics**. (Top) ANOSIM values obtained by subsetting and dividing the data into different groups. (Bottom) Number of communities for starting and final classes.

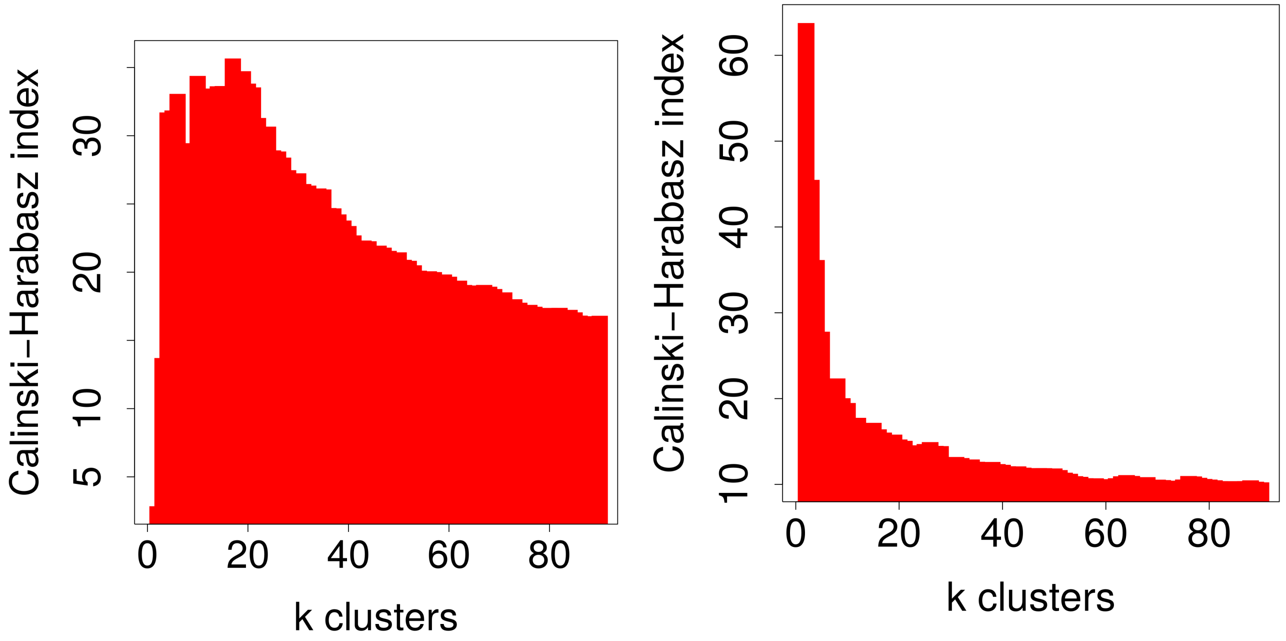

Figure 4: **Determination of number of community classes.** Communities were clustered according to their Jensen-Shannon distance using a partition-around-medoids clustering for an increasing number of clusters $k$ (x-axis), and the Calinski-Harabasz index computed to estimate the optimal clustering. (Left) Starting communities have an absolute maximum when communities are classified in 17 clusters. In previous work, we worked with the same communities defined by Operational Taxonomic Units finding a maximum at 6 clusters, which means that ASVs allow us to identify a higher variability. For consistency with previous work, here we also work with the classification in 6 clusters, which corresponds to the second maximum. (Right) Final communities have an absolute maximum when communities are classified in 2 clusters. Results are for replicate 1. The other three replicates are similar.

#### Statistical analysis of metagenomics predictions

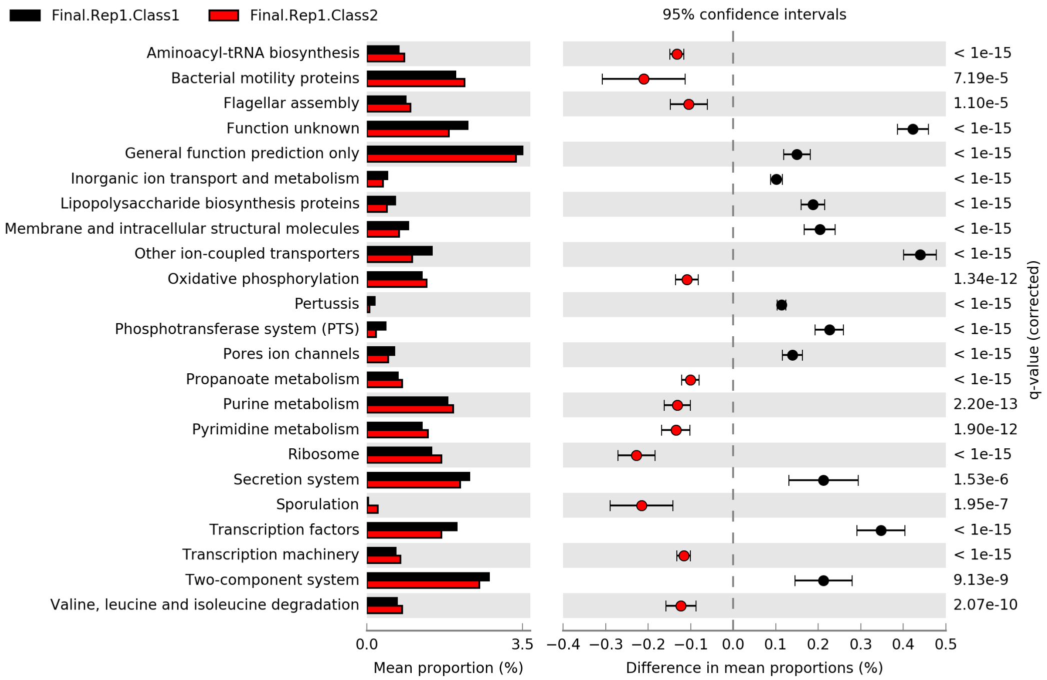

Figure 5: **Example of statistical test for the analysis of metagenome pathways.** Rows indicate KEGG pathways found significant in the comparison between class 1 and class 2 of FC. The first column is the mean proportion of each class, the second column the difference in mean proportions and the third column the Benjamini-Hochberg corrected p-value (termed q-value) of a Welch's test. Only pathways with a difference larger than 0.1 are shown. Significant pathways found in the comparison between class 1 and class 2 for each replicate were used to create the heatmap shown in the Main Text.

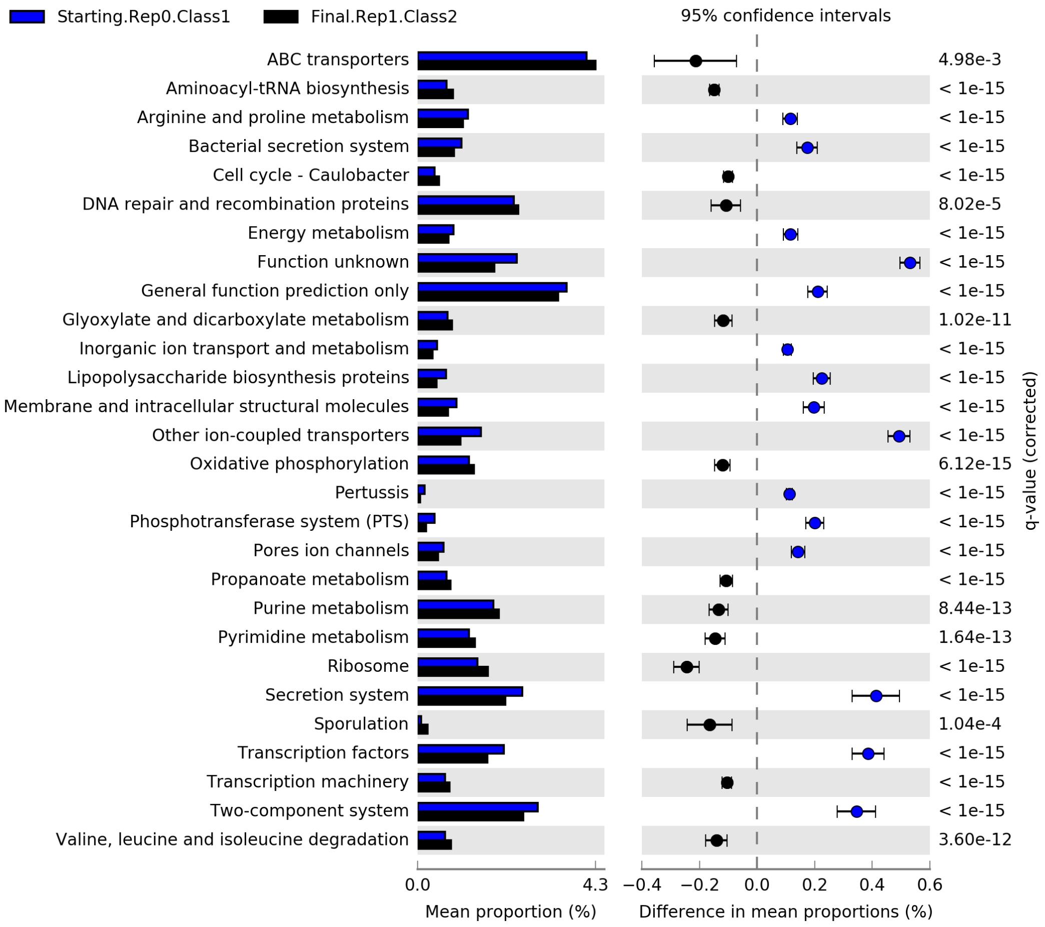

Figure 6: **Example of statistical test for the analysis of metagenome pathways.** Rows indicate KEGG pathways found significant in the comparison between SC class 1 and FC class 2. The first column is the mean proportion of each class, the second column the difference in mean proportions and the third column the Benjamini-Hochberg corrected p-value (termed q-value) of a Welch's test. Only pathways with a difference larger than 0.1 are shown. Significant pathways found in these comparisons were used to create the heatmap shown in Suppl. Fig 7.

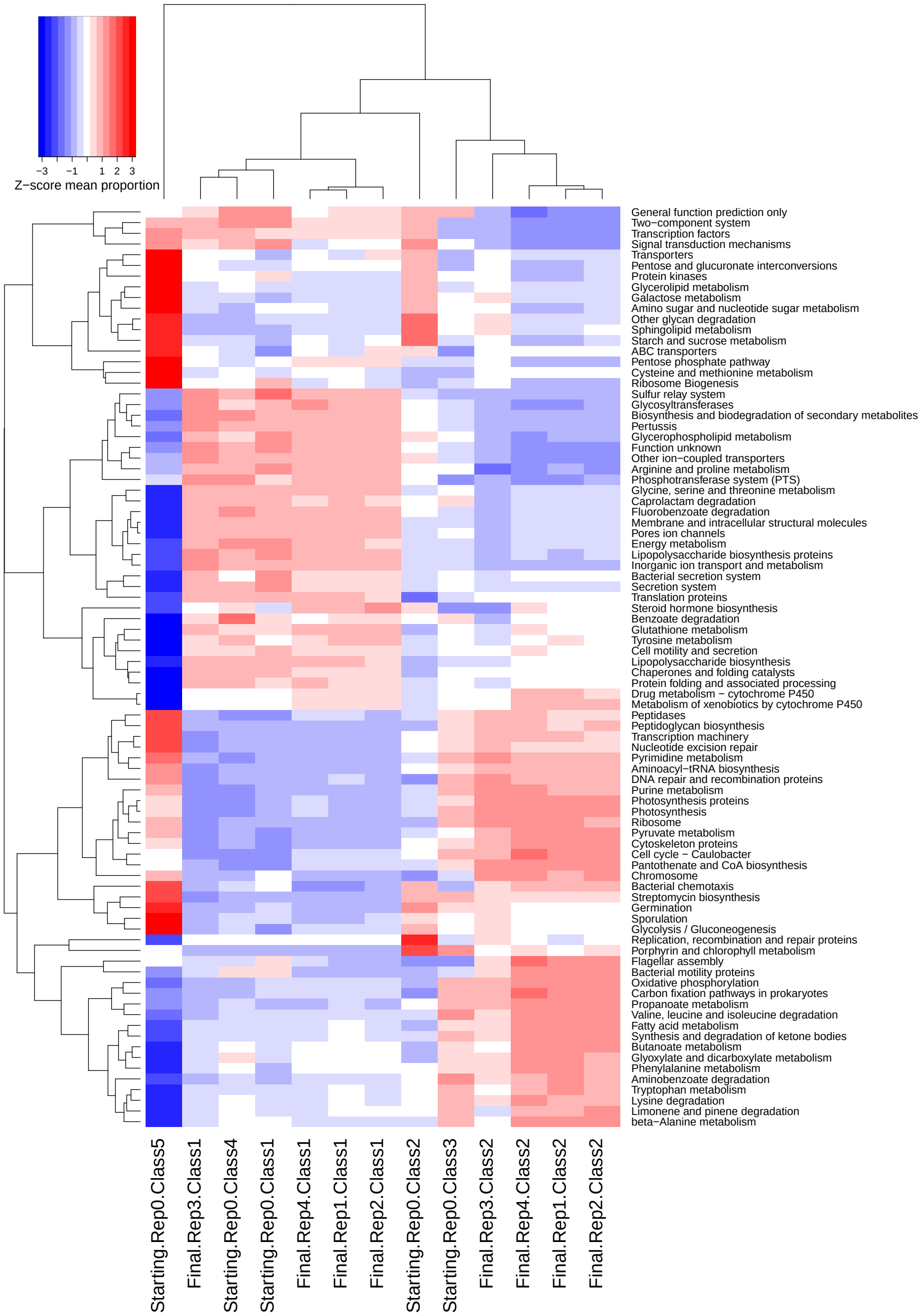

Figure 7: **Metagenomic convergence.** Z-score of the difference in mean proportion of genes clustered in KEGG metabolic pathways between starting- and final community classes. The scaling of the Z-score was computed for each pathway (i.e. scaled by rows), and only pathways showing a significant difference are shown (Welch test corrected for multiple testing). Clustering of classes is similar to the compositional clustering shown in Fig. 10, with starting classes 1 and 4 joining final class 1, while final class 2 cluster independently. Starting community classes 2 and 3 appear intermediate between both. Starting class 5 seems to be an outlier, with very significant pathways. This means that its communities are functionally very similar but note, however, that this class hosts the lowest number of communities (only 10).

#### Landscape transformation and equivalent classes

Since the four replicates of final communities had their optimal classification for two clusters, we asked if these classes were equivalent across replicates. To answer this question, we computed all-against-all Jensen-Shannon distances between all samples, i.e. including starting communities and the four replicates of final communities. We then computed the mean distance within each class and between classes, which were determined independently for each dataset. We found that the first (second) class of final communities were more similar among themselves than they were with respect to the second (first) class found in their own replicate, suggesting that the classifications were equivalent (Suppl. Fig. 10). Interestingly, starting community classes 1 and 4 (containing the largest set of communities) were clustered with final community class 1 and have a high similarity, suggesting that it is a large stable attractor. On the other hand, starting community classes 2, 3 and 5 clustered with the final community class 2, but their mean similarity is much lower, suggesting that the new attractor represented by final community class 2 demands a more marked transformation in the starting community composition.

The transformation of the compositional landscape is illustrated in Suppl. Fig. 8, together with other potential scenarios. The dependence of the trajectories from starting classes membership is shown in Fig. 9.

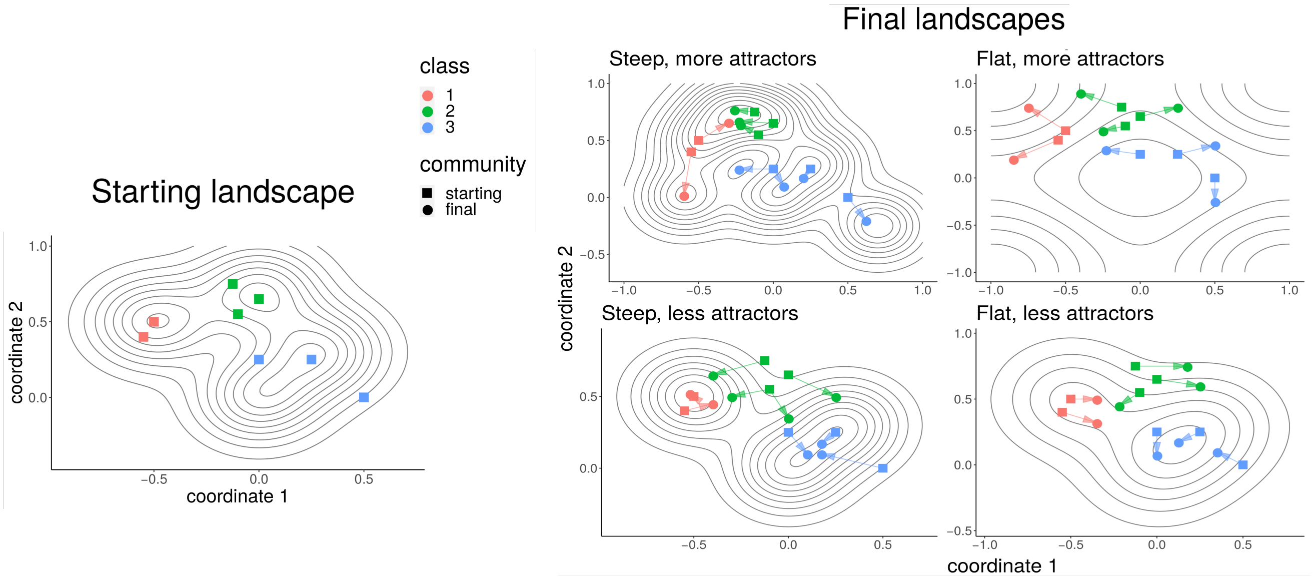

Figure 8: **Transformation of the compositional landscape.** Illustration showing an imaginary transformation in which a starting landscape can be transformed into one of four possible hypothetical final landscapes. (Left) Illustration of the compositional landscape from the perspective of the starting communities. Three attractors corresponding to three community classes are shown, with squares representing the starting communities. (Right) Four hypothetical final landscapes showing an increase/decrease in the number of attractors and the slope of the landscape with respect to the initial landscape. The trajectories of the initial communities depend on the initial positions relative to the final landscape. Our results are described by a scenario in which the number of attractors is reduced, with one of them (Final Class 1) being steep and Final Class 2 possibly flat.

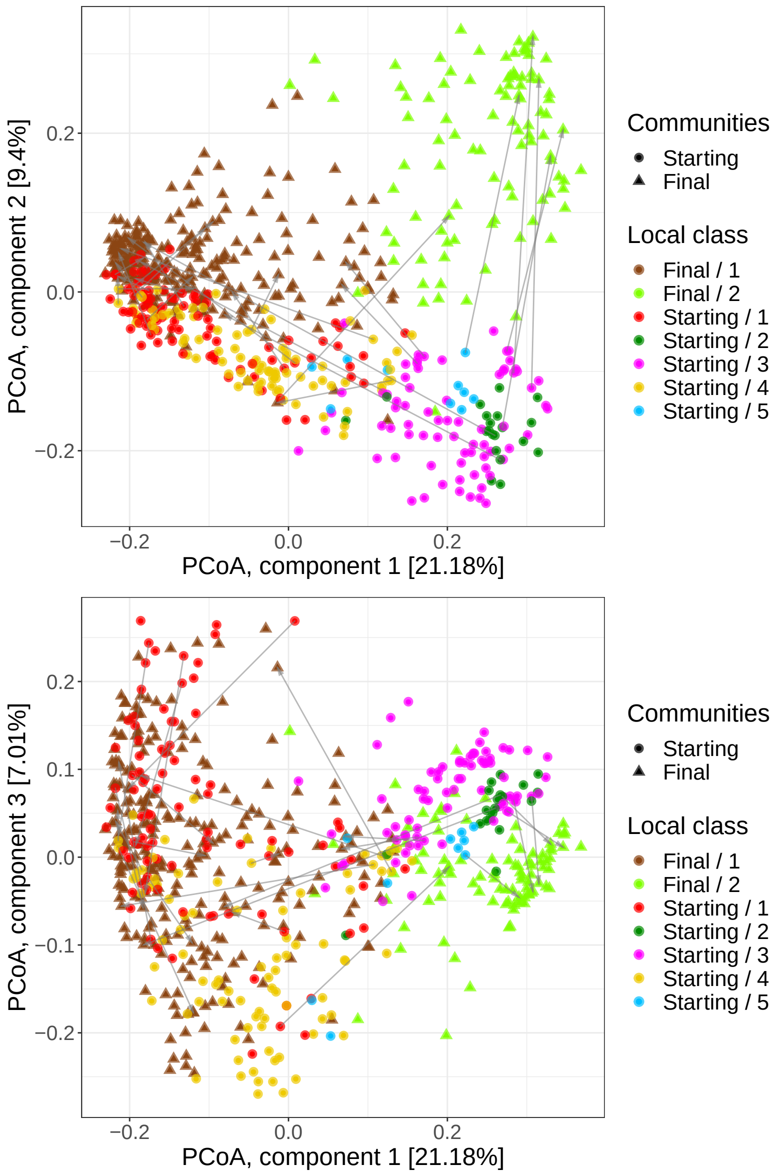

Figure 9: **Correspondence between starting and final classes.** PCoA of the starting- and final communities for one of the replicates showing the correspondence between the starting community and the two final community classes for components 1 and 2 (top) and 1 and 3 (bottom). We observed that the number of attractors was reduced and that the topography likely flattened because the points became more dispersed, especially for final class 2. A random subset (10%) of the trajectories are shown using arrows.

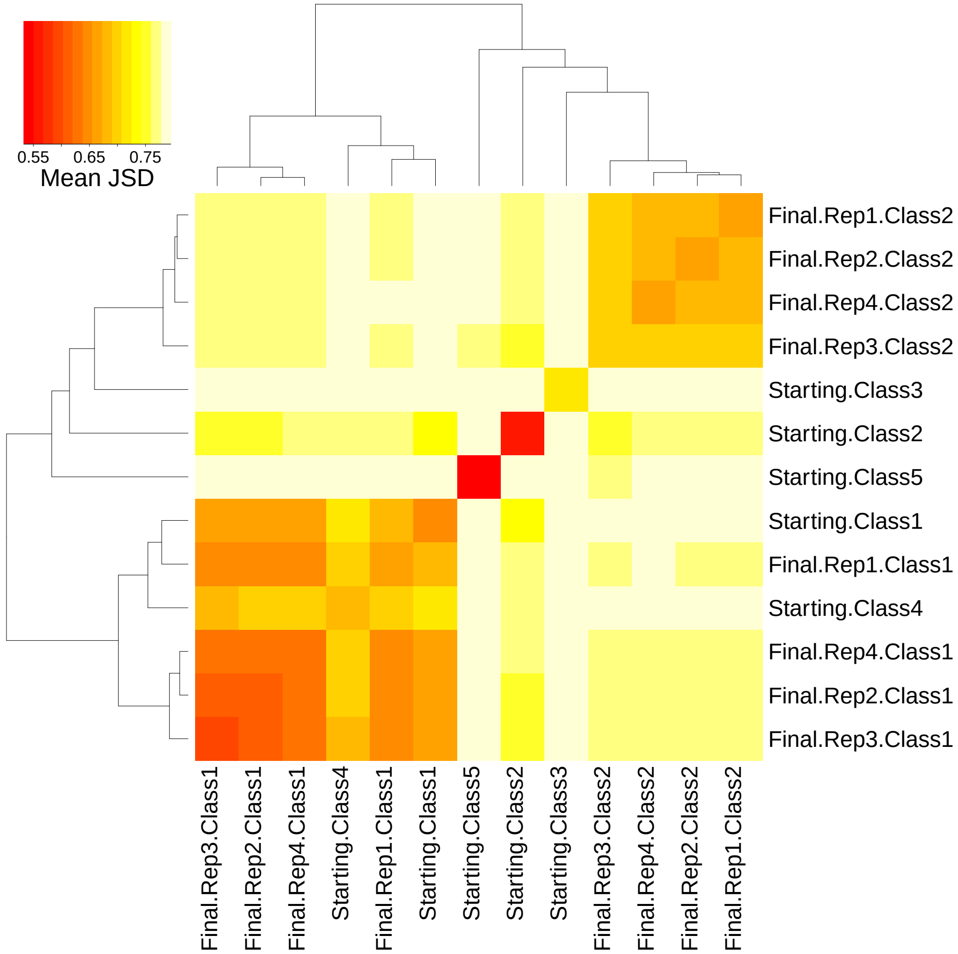

Figure 10: **Similarity between community classes.** Heatmap showing the mean Jensen-Shannon diversity between communities belonging to different classes (starting communities, labelled starting, and the four replicates of the final communities, labelled final). Starting community classes 1 and 4 cluster with the four replicates of final community classes 1. Final community class 2 also cluster together, with starting community class 2 being the more similar one. Starting community class 3 and 5 are the most dissimilar to any other class.
